# Phylogeny is not taxonomy: lessons on higher classification from turtle and whale barnacles (Cirripedia: Coronuloidea)

**DOI:** 10.64898/2026.09.05.749650

**Authors:** Ryota Hayashi, Alberto Collareta

**Affiliations:** Center for Advanced Research, Research & Development Center, Nippon Koei Co., Ltd., 2304 Inarihara Tsukuba, Ibaraki, 300-1259, Japan; Dipartimento di Scienze della Terra, Università di Pisa, via S. Maria 53, 56126 Pisa, Italy; Museo di Storia Naturale, Università di Pisa, Via Roma 79, 56011 Calci, Italy

**Author notes:** Corresponding author: Ryota Hayashi, Research & Development Center, Nippon Koei Co., Ltd., 2304 Inarihara Tsukuba, Ibaraki, 300-1259, Japan.

**Keywords:** classification, monophyly, diagnosability, stability, utility

## Abstract

Phylogeny and taxonomy are related but not identical: phylogeny estimates common ancestry, whereas taxonomy must also provide diagnosable, stable, informative, and useful classifications. We examine this distinction using turtle and whale barnacles of the superfamily Coronuloidea, a group with an extensive fossil record, marked ecological specialization, a long history of morphology-based classification, and a recent topology-driven reclassification that merged recognized higher taxa to preserve monophyly. These changes reduced diagnosability and biological coherence, weakened historical continuity, and relied on placements insufficiently tested against morphological and phylogenetic evidence. Our reanalysis of published molecular datasets shows that critical relationships are sensitive to marker composition and taxon sampling. Together with morphological considerations on extant and fossil forms, these results indicate that topology alone is insufficient for coherent higher classification. We argue that higher taxa should be evaluated using complementary criteria—diagnosability, stability, utility, and monophyly—rather than by mechanically translating a preferred tree into Linnaean ranks.

“All true classification is genealogical; that community of descent is the hidden bond which naturalists have been unconsciously seeking.” —Darwin, 1859

## Introduction

The relationship between phylogeny and taxonomy has been debated for more than half a century. Although the two are intimately related, they are not the same enterprise. A phylogeny estimates branching history, whereas a classification organizes biological information into ranked units that can be stored, communicated, retrieved, and used comparatively. The enduring challenge is therefore not to choose between phylogeny and taxonomy, but to determine how they should relate to each other in systematic practice.

This distinction was articulated clearly in the classical literature. Bigelow (1958) argued that classification based on overall similarity and classification based on recentness of common ancestry are not equivalent, because evolution is change rather than time alone. Bock (1973) later defended evolutionary classification as a system intended to summarize both common descent and biologically meaningful similarity. Ashlock (1979) also emphasized that higher classification should maximize utility rather than merely mirror one formal criterion. Brummitt (1997) further sharpened this distinction by arguing that classification and phylogeny have different functions and should be allowed to coexist, while Franz (2005) diagnosed a growing “phylogeny/classification gap” as a symptom of modern systematics drifting away from classification as a communicative scientific practice.

These old debates remain highly relevant today. Although shared ancestry is rightly treated as a prime source of systematic information, many authors have pointed out that monophyly alone does not settle the problem of how ranked taxa should be delimited and named. Vences *et al*. (2013) explicitly proposed naming criteria that include not only monophyly, but also clade stability and phenotypic diagnosability. Wilkerson *et al*. (2015) argued that a stable Linnaean classification must balance utility with current evolutionary knowledge rather than track every poorly supported rearrangement. More recently, Kuntner *et al*. (2023) argued that ranked taxa above the species level should maximize information content, diagnosability, and utility, and that classifications should not automatically collapse historically meaningful taxa into oversized, weakly informative units simply because new phylogenetic results reveal asymmetry in diversity.

The problem is especially acute at higher taxonomic ranks. Families, superfamilies, and orders are not merely nested branches of a cladogram; they are comparative units that summarize broad morphological organization, ecological shifts, fossil information, and historically accumulated biological knowledge. As a result, a ranked taxon may be technically monophyletic yet still taxonomically poor if it is weakly diagnosable, biologically diffuse, unstable, or otherwise uninformative. Conversely, a classification may proves most useful if it preserves historically meaningful and diagnostically coherent taxa while remaining broadly consistent with phylogenetic evidence. In this sense, higher classification cannot be reduced to the mechanical translation of a preferred topology into Linnaean ranks.

This problem is illustrated particularly clearly by the turtle and whale barnacles traditionally classified in the superfamily Coronuloidea. Historically, coronuloid systematics was shaped by morphology, functional adaptation, and host association, in a tradition extending from Charles Darwin’s monumental work on cirripedes through later, 20th century classifications (Darwin 1854, Pilsbry 1916, Newman and Ross 1976, Monroe 1981, Newman 1996), as summarized in Figure 1. Family boundaries were drawn from shell shape, wall structure, patterns of opercular plate reduction, and the contrast between rock-dwelling barnacles and epibiotic barnacles specialized for life on sea turtles, cetaceans, sirenians, and other living hosts (Hayashi 2013). Even where these morphology- based systems disagreed, they remained biologically interpretable. In its traditional sense, Coronuloidea thus refers to a striking adaptive radiation of obligate or near-obligate epibiotic barnacles.

**Figure 1.**
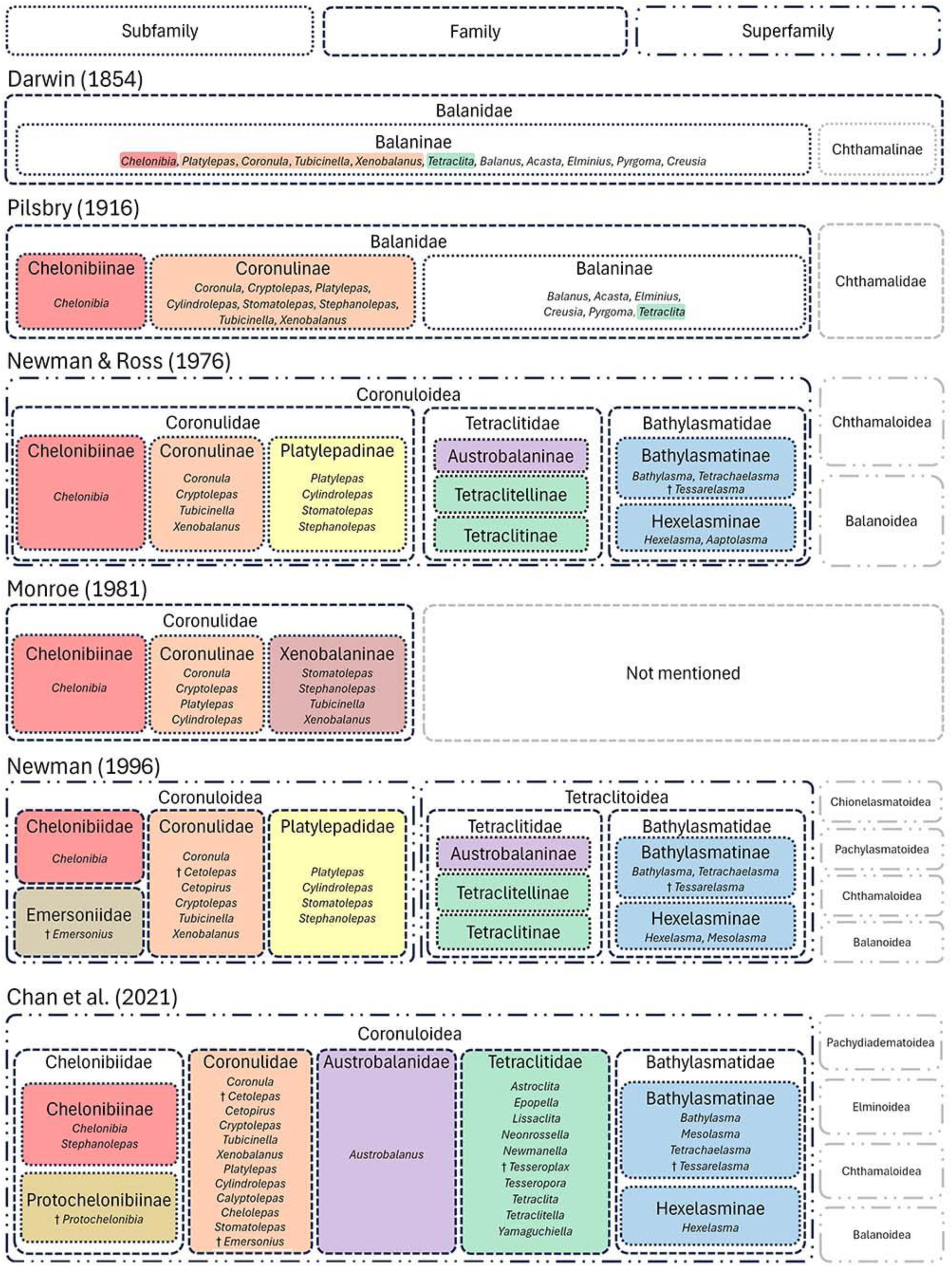
Historical changes in the higher-level classification of Coronuloidea and related balanomorph barnacles from Darwin (1854) through major subsequent classifications, culminating in Chan et al. (2021). Each column summarizes a representative classification scheme adopted by Darwin (1854), Pilsbry (1916), Newman & Ross (1976), Monroe (1981), Newman (1996), and Chan et al. (2021).

Broader molecular phylogenetic studies subsequently expanded the picture (e.g., Pérez-Losada *et al*. 2004, 2008), but they included only a limited coronuloid sample, leaving the superfamily’s internal relationships largely unevaluated. Building on that context, Hayashi *et al*. (2013) focused specifically on turtle and whale barnacles, and showed that several traditional family-level circumscriptions were not replicated phylogenetically. In particular, *Stephanolepas* was recovered as sister to *Chelonibia*, Platylepadidae proved polyphyletic, and *Cylindrolepas* itself was not found to be monophyletic.

These results were important because they showed that several traditional taxonomic concepts within the superfamily required re-examination. Hayashi *et al*. (2013), however, did not treat the lack of evidence for monophyly as a sufficient reason to establish a new family-level classification; the study left rank assignment open rather than translating each inferred relationship directly into revised Linnaean ranks.

The higher classification proposed by Chan *et al*. (2021) for barnacles brings this problem into sharp focus. In that ambitious synthesis, the long-established superfamily Tetraclitoidea was abandoned in favor of a very broad concept of Coronuloidea that also encompasses Chelonibiidae, Coronulidae, Tetraclitidae, Bathylasmatidae, and Austrobalanidae, with Platylepadidae being dissolved into Coronulidae. In effect, this treatment translated a limited set of molecular topologies directly into a revised higher classification. However, the resulting taxa were diagnosed in ways that appear largely post hoc and weakly informative. As a consequence of this, traditional names were broadened, and their morphological coherence and comparative significance were substantially reduced.

Here, we reassess the higher classification of Coronuloidea by integrating reanalyses of published molecular datasets with comparative observations on shell architecture, opercular morphology, ecology, and fossil evidence. We use this case to evaluate whether the topology-driven expansion of Coronuloidea and dissolution of Platylepadidae proposed by Chan *et al*. (2021) improve classificatory coherence, or instead reduce diagnosability, stability, and taxonomic utility. Our aim is not to deny the importance of molecular phylogeny, nor to defend every traditional family-level arrangement as unchanged, but to clarify how phylogenetic evidence should be translated into ranked higher classification.

## Materials and Methods

### Materials examined and morphological observations

Specimens representing the principal coronuloid and tetraclitoid lineages discussed in this study were examined comparatively to document major differences in shell architecture and opercular morphology. For each taxon, the shell was photographed in apical, basal, and lateral views, and the scutum and tergum were examined both internally and externally. Representative members of Coronuloidea and other balanomorph barnacles are shown in Figures 2 and 3.

**Figure 2.**
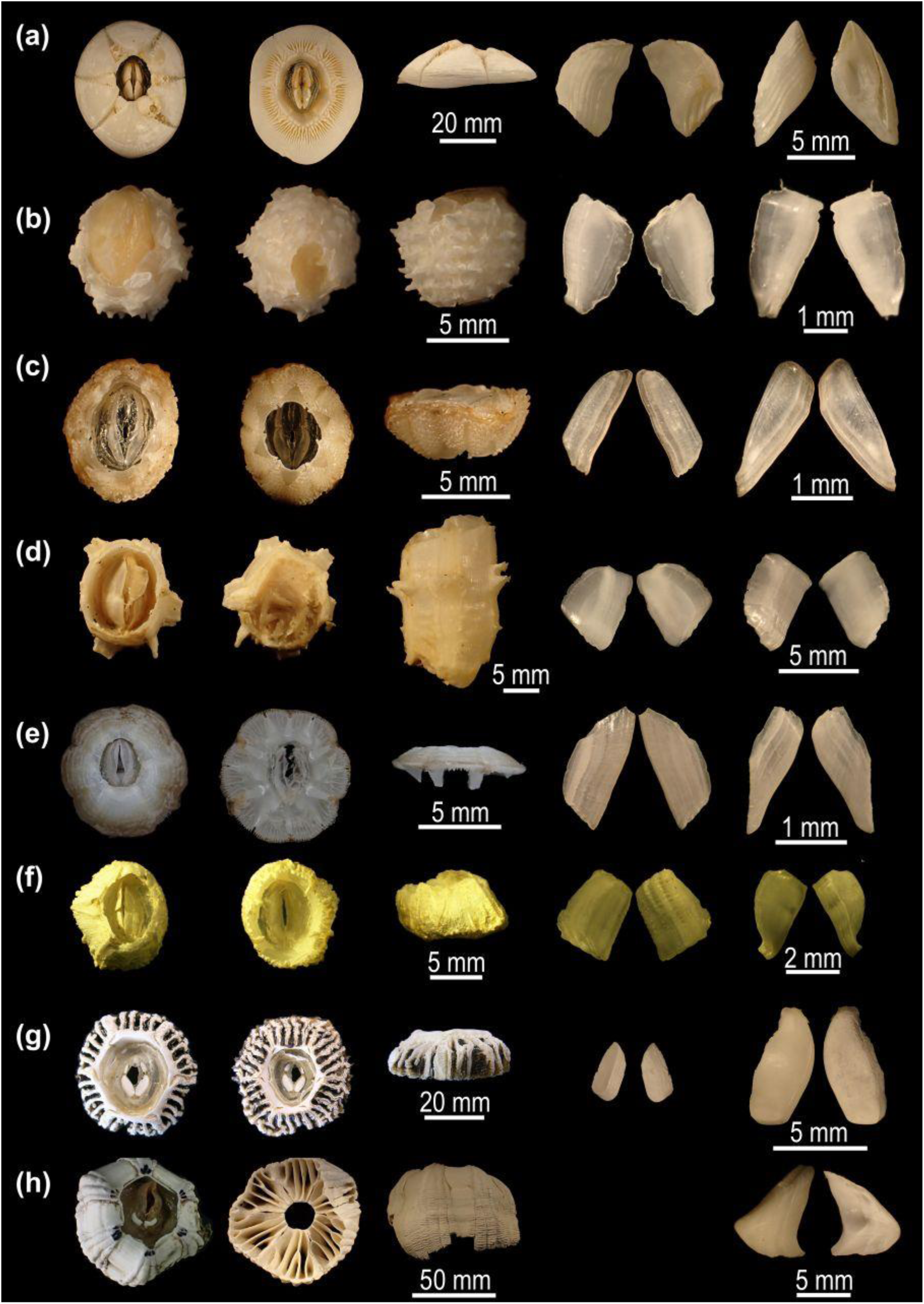
Barnacles of Coronuloidea. From left to right: apical shell view, basal shell view, lateral shell view, outer and inner surfaces of tergum and scutum. (a) *Chelonibia testudinaria*; (b) *Stephanolepas muricata*; (c) *Stomatolepas praegustator*; (d) *Tubicinella cheloniae*; (e) *Platylepas hexastylos*; (f) *Cylindrolepas darwiniana*; (g) *Cryptolepas rhachianecti*; (h) *Coronula diadema*.

**Figure 3.**
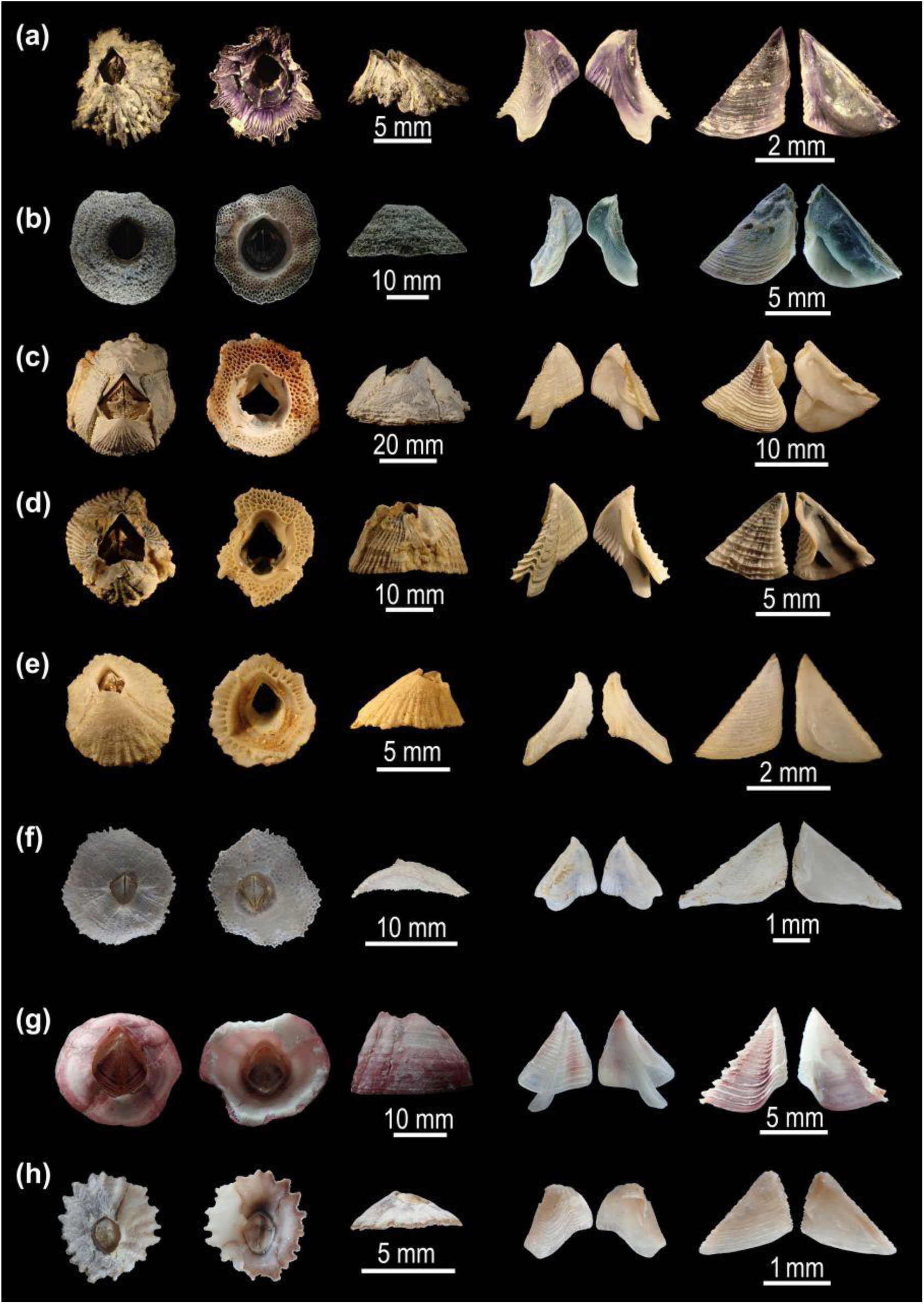
Barnacles of Tetraclitoidea and other superfamilies. From left to right: apical shell view, basal shell view, lateral shell view, outer and inner surfaces of tergum and scutum. (a) *Austrobalanus imperator*; (b) *Tetraclita japonica*; (c) *Newmanella spinosus*; (d) *Yamaguchiella coeruleascens*; (e) *Tesseropora alba*; (f) *Tetraclitella divisa*. A member of Balanoidea, *Megabalanus rosa*, is shown in (g), whereas a member of Chthamaloidea, *Chthamalus moro*, is shown in (h).

### Phylogenetic analyses

The molecular dataset analyzed herein was assembled mainly from sequences previously published by Hayashi *et al*. (2013) and Tsang *et al*. (2015), with additional sequences obtained from Spears *et al*. (1994), Pérez-Losada *et al*. (2004, 2008), Simon-Blecher *et al*. (2007), Tsang *et al*. (2012), and Cheang *et al*. (2013). Taxa included in the present study and their GenBank accession numbers are listed in Supplementary Table S1. Each marker was aligned separately using MAFFT (Katoh *et al*. 2019), and the resulting alignments were subsequently inspected and manually adjusted where necessary to improve positional homology. Phylogenetic trees were inferred under the maximum-likelihood criterion using IQ-TREE (Trifinopoulos *et al*. 2016).

To isolate the effect of locus sampling while holding taxon sampling constant, we analyzed the same dataset for 28 operational taxonomic units (OTUs) under two marker combinations: a four-marker dataset comprising 12S, 16S, 18S, and H3, and a five-marker dataset comprising 12S, 16S, 18S, 28S, and H3. The second dataset for 45 OTUs included four markers (12S, 16S, 18S and H3) to enable broader taxon sampling.

Previous molecular studies suggested that *Chelonibia testudinaria*, *C. patula*, and *C. manati* are not genetically distinguishable, thus possibly representing a single species (Cheang *et al*. 2013, Zardus *et al*. 2014). However, for consistency with the sampling design of Hayashi *et al*. (2013), here we treat them as separate OTUs. We also follow Hayashi (2012) in treating *Tubicinella cheloniae* within *Tubicinella* rather than within *Chelolepas*.

## Results

### Comparative morphology of coronulid barnacles and other balanomorph barnacles

Comparative examination of the taxa illustrated in Figures 2 and 3 revealed marked differences in shell architecture and opercular morphology between the classical coronuloids and other balanomorphs, including the tetraclitoids. The coronuloids taxa examined here exhibit a broad spectrum of shell modifications associated with epibiotic life, including a low, encrusting shell in *Chelonibia testudinaria*; a deep, host-penetrating shell in *Stephanolepas muricata*; an even more tube-like shell in *Tubicinella cheloniae*, a flattened shell in *Platylepas hexastylos*, and a large, deeply folded shell in *Coronula diadema* (Fig. 2). Their opercular plates are variably reduced, simplified, or otherwise modified. By contrast, *Austrobalanus imperator*, the tetraclitids, and other non-coronuloid balanomorphs examined for comparison retain more generalized shell forms and well-developed scuta and terga (Fig. 3). These observations document a clear morphological contrast between the radiation of the turtle and whale barnacles and the overall more conservative tetraclitid– bathylasmatid–austrobalanid assemblage.

### Phylogenetic sensitivity and locus discordance

The maximum-likelihood analyses of the two concatenated datasets recovered broadly similar higher-level structure, in that Coronuloidea and Tetraclitoidea were separated as major lineages, but several internal relationships differed between the 4- and 5-marker trees (Fig. 4).

**Figure 4.**
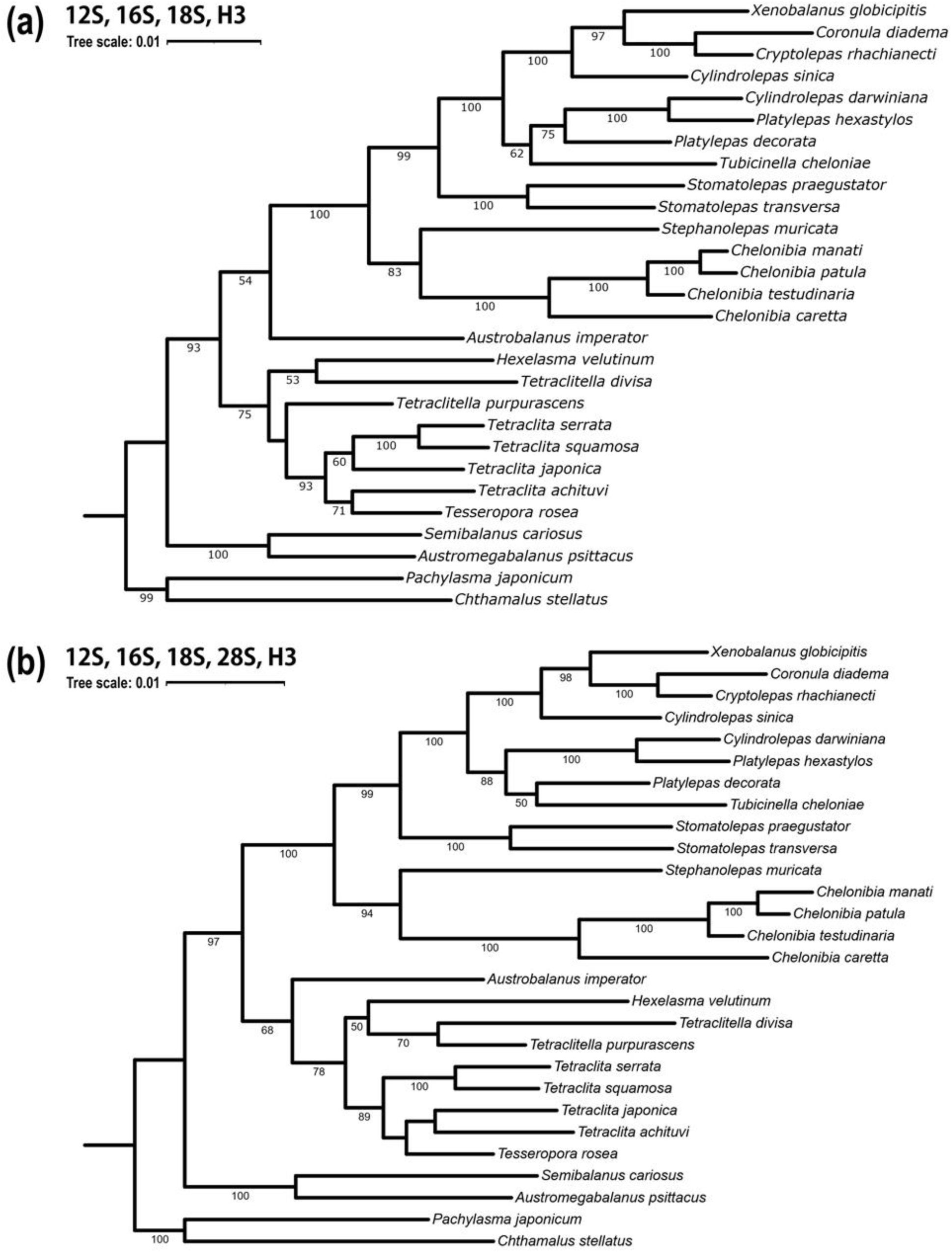
Comparison of concatenated maximum-likelihood phylogenies inferred from the same 28- OTU dataset under two different marker combinations. (a) Tree inferred from four markers (12S, 16S, 18S, and H3). (b) Tree inferred from five markers (12S, 16S, 18S, 28S, and H3). Numbers at nodes indicate bootstrap support values.

In the 4-marker tree (Fig. 4a), *Platylepas decorata* appeared as sister to the clade comprising *P. hexastylos* and *Cylindrolepas darwiniana*. By contrast, in the 5-marker tree (Fig. 4b), *P. decorata* and *Tubicinella cheloniae* formed a clade.

The phylogenetic position of *Austrobalanus* also differed between the two analyses. In the 4-marker tree, *Austrobalanus* was placed as sister to the classical Coronuloidea, rendering the classical Tetraclitoidea paraphyletic (Fig. 4a). In contrast, in the 5-marker tree, it was recovered as the earliest-diverging lineage within the classical Tetraclitoidea (Fig. 4b). In addition, *Austrobalanus* was recovered in different placements depending on the marker: in the 12S tree, it appeared as sister to a clade comprising *Bathylasma*, *Hexelasma*, and the tetraclitids (Fig. 5a); in the 16S tree, it was placed within the coronuloid assemblage, as sister to *Stephanolepas muricata* (Fig. 5b); in the 18S tree, it was embedded within the broader tetraclitoid assemblage (Fig. 5c); in the 28S tree, it formed an isolated branch sister to a reduced tetraclitoid clade (Fig. 5d); and in the H3 tree, it clustered with *Tetraclita rubescens* (Fig. 5e). Thus, the phylogenetic position of *Austrobalanus* was highly unstable across markers.

**Figure 5.**
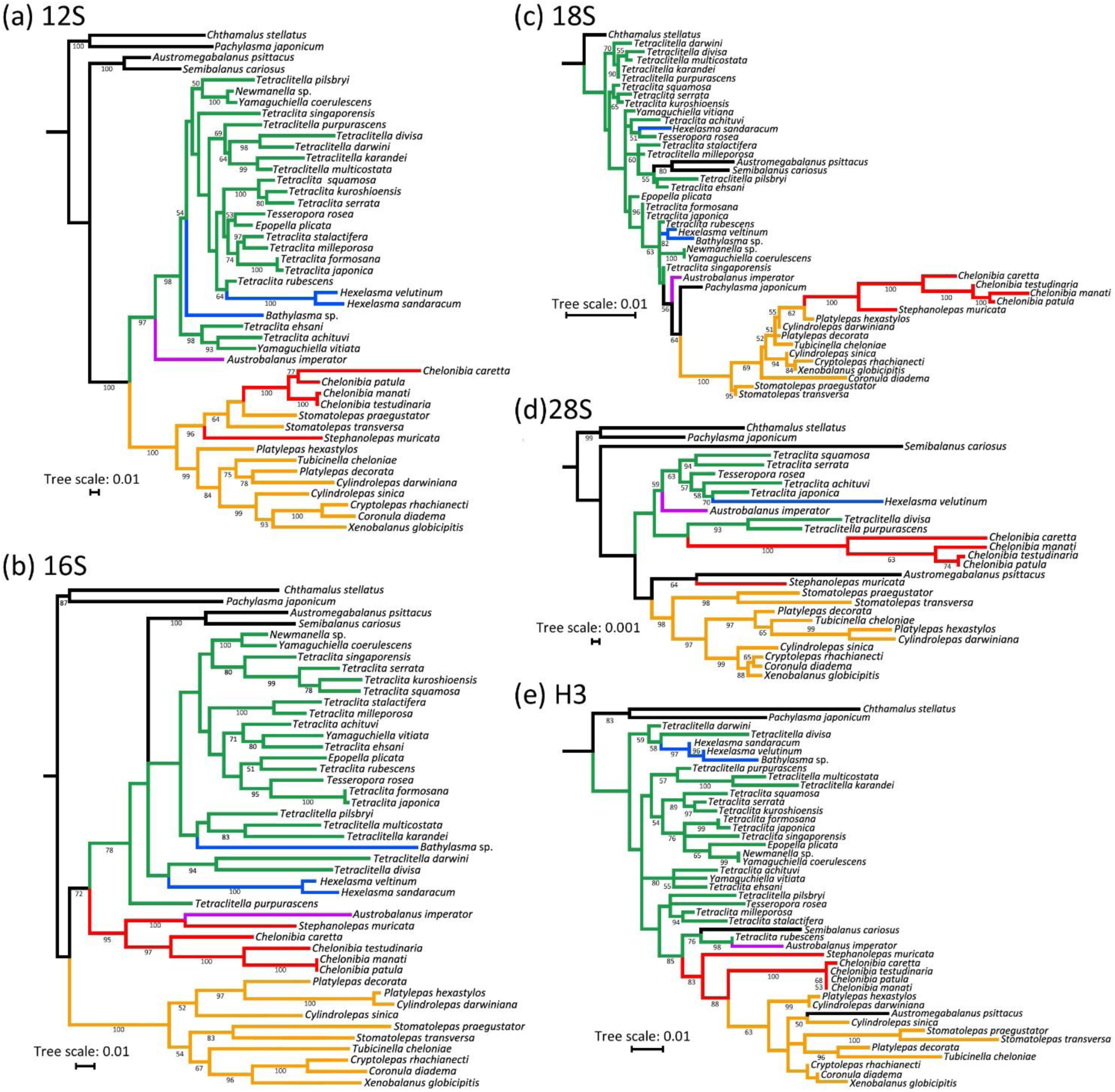
Phylograms inferred from five molecular markers: (a) 12S, (b) 16S, (c) 18S, (d) 28S, and (e) H3. Branches are color-coded according to the family-level classification of Chan et al. (2021): Chelonibiidae, Coronulidae, Tetraclitidae, Bathylasmatidae, and Austrobalanidae. Taxa outside these focal families are shown in black. Scale bars indicate substitutions per site.

A further difference concerned the placement of *Hexelasma*. In the 4-marker tree, *Hexelasma velutinum* was recovered clustering with *Tetraclitella divisa*, but apart from *Tetraclitella purpurascens* (Fig. 4a and S1). In the 5-marker tree, however, *H. velutinum* formed a clade with *Tetraclitella* spp. (Fig. 4b), indicating instability in the relationship between these taxa under the two sampling schemes. The monophyly of *Bathylasma*/*Hexelasma* was likewise inconsistent among loci. In the 12S, 16S, and 18S trees, *Bathylasma* did not group with *Hexelasma* (Fig. 5a-c). In the 28S tree, *Bathylasma* was absent, so that the monophyly of *Bathylasma* + *Hexelasma* could not be evaluated (Fig. 5d). In contrast, the H3 tree recovered *Bathylasma* in a clade with both *Hexelasma* spp. (Fig. 5e).

The position of *Stephanolepas* also differed among loci, yet it was consistently associated with coronuloid taxa. In the 12S tree, *Stephanolepas* was found at the base of a clade including *Stomatolepas* and *Chelonibia* (Fig. 5a). In the 16S tree, it was recovered with *Austrobalanus*, and the pair was sister to *Chelonibia* (Fig. 5b). In the 18S tree, *Stephanolepas* alone was sister to the *Chelonibia* clade (Fig. 5c). In the 28S tree, *Stephanolepas* formed a clade with *Austromegabalanus*, whereas *Chelonibia* was nested within a disrupted tetraclitoid assemblage including *Tetraclita*, *Tetraclitella*, *Tesseropora*, *Hexelasma*, and *Austrobalanus* (Fig. 5d). In the H3 tree, *Stephanolepas* was positioned adjacent to the *Chelonibia* clade (Fig. 5e).

Four loci yielded especially conspicuous placements of non-coronuloid, non-tetraclitoid taxa within Tetraclitoidea or Coronuloidea. The 16S tree placed *Austromegabalanus* and *Semibalanus* within the broad tetraclitoid assemblage (Fig. 5b). The same was true for the 18S tree, where *Pachylasma* was further recovered as sister to the classical Coronuloidea (Fig. 5c). In the 28S tree, *Austromegabalanus* was recovered within the coronuloid assemblage (Fig. 5d). In the H3 tree, *Austromegabalanus* was again embedded within the classical Coronuloidea, with *Semibalanus* being recovered as sister to a clade including *T. rubescens* and *Austrobalanus* (Fig. 5e). These results indicate substantial discordance among loci in the placement of OTUs belonging to major higher taxa.

Overall, the concatenated and single-marker analyses indicate that several traditional higher taxa are not consistently recovered as monophyletic, and that the positions of key lineages, especially *Austrobalanus* and *Stephanolepas*, are highly sensitive to marker choice. Comparison with the broader 45-OTU four-marker analysis further indicates that expanded taxon sampling mainly affects relationships within the tetraclitid–bathylasmatid assemblage, while leaving the position of *Austrobalanus* relative to the classical Coronuloidea largely unchanged under the four-marker framework (Fig. S1).

## Discussion

### When topology drives classification: the case of turtle and whale barnacles

The present analyses indicate that the higher-level placement of *Austrobalanus imperator* is highly sensitive to several analytical dimensions. First, comparison of the two 28-OTU concatenated trees shows a clear effect of marker composition: *Austrobalanus* was recovered as sister to the classical Coronuloidea in the four-marker analysis, but as the earliest-diverging lineage within the classical Tetraclitoidea when 28S was added (Fig. 4). Second, single-marker analyses revealed an even greater locus-specific discordance, placing *Austrobalanus* in different coronuloid, tetraclitoid, or isolated positions depending on the marker analysed (Fig. 5). Third, comparison with the broader 45-OTU four-marker analysis suggests that expanded taxon sampling modifies relationships within the tetraclitid–bathylasmatid assemblage, even though it does not substantially alter the placement of *Austrobalanus* relative to the classical Coronuloidea under the four-marker framework. Together, these results indicate that the apparent position of *Austrobalanus* depends on marker composition, marker-specific signal, and the sampling context around the traditional Coronuloidea–Tetraclitoidea boundary.

This interpretation also helps explain the difference between the present analyses and that by Hayashi *et al*. (2013). That study recovered *Austrobalanus* near the base of Coronuloidea in a five-marker analysis, but included only sparse sampling of Tetraclitoidea. By contrast, the urrent work includes a broader sampe of the tetraclitid–bathylasmatid–austrobalanid taxa and recovers different placements of *Austrobalanus* depending on marker composition and locus-specific signals. This discrepancy are not be attributed to a single factor, such as taxon sampling alone or marker composition alone. Rather, it reinforces the conclusion that the placement of *Austrobalanus* remains provisional and should not be used as a stable basis for expanding Coronuloidea. The taxonomic question is therefore not simply which topology is preferred, but what classificatory information is gained or lost when that topology is translated into ranks.

This instability matters because the broad concept of Coronuloidea proposed by Chan *et al*. (2021) does more than recognizing a molecularly inferred assemblage —it transforms the whole biological meaning of a long-established superfamily. By absorbing austrobalanids, bathylasmatids, and tetraclitids, Coronuloidea becomes more inclusive, but also more morphologically diffuse, ecologically heterogeneous, and historically detached from its traditional usage. *Austrobalanus* and the tetraclitids are primarily rock-dwelling intertidal barnacles, and bathylasmatids are predominantly deep-sea forms, whereas the classical coronuloids comprise a specialized radiation centered on epibiosis on marine vertebrates and other living hosts. The contrast is also morphological: classical coronuloids—the turtle and whale barnacles—are characterized by variably reduced (or even absent) opercular plates, whereas tetraclitids and kin retain well-developed scuta and terga typical of generalized balanomorph barnacles (Figs. 2 and 3).

Lumping these lineages into a single superfamily may reduce conflict with one preferred topology, but also decreases coherence in anatomy, ecology, and biological interpretation. The rank no longer summarizes a distinctive adaptive radiation, instead becoming a broad label for morphologically and ecologically disparate forms. This is not merely a matter of taxonomic taste. A higher taxon can be technically monophyletic yet still be taxonomically poor if it is too diffuse to diagnose, too broad to communicate efficiently, and/or too weakly connected to the comparative inferences historically associated with that name. In this respect, Coronuloidea sensu Chan *et al*. (2021) is better understood as a topology-driven circumscription than as a morphology-led taxonomic redefinition.

The same problem appears at the family level. Chelonibiidae sensu Chan *et al*. (2021) includes both *Chelonibia* and *Stephanolepas*—a grouping that conflicts with fundamental morphological differences in shell architecture and opercular structure: *Chelonibia* shows a generalized, encrusting form (Fig. 2a), whereas *Stephanolepas* possesses a deep, host-penetrating shell with an inverted profile and a simplified tergum (Fig. 2b). The familial placement by Chan *et al*. (2021) of fossil-only taxa such as †*Emersonius* is also difficult to justify, as it builds upon a strictly molecular scaffold without explicit morphological argumentation (Perreault *et al*. 2025). Such placement may appear precise, yet it rests on indirect, largely passive inference—one that comes at the cost of weakening the role of the fossil record as an independent source of information. Similarly, the dissolution of Platylepadidae into Coronulidae removes a historically meaningful family without providing a correspondingly stronger diagnosis for the surviving, inflated family. Coronulidae sensu Chan *et al*. (2021) therefore functions as an accommodative family concept: it absorbs heterogeneous lineages that no longer fit older boundaries, but does so without gaining a clearer diagnostic or comparative meaning.

### Toward a more informative classification: four criteria for taxonomy

The distinction between a taxon and the characters used to recognize it predates evolutionary systematics. Linnaeus famously expressed the point as “Scias Characterem non constituere Genus, sed Genus Characterem”, which translates as “Know that the character does not constitute the genus, but the genus the character” (Linnaeus 1751). Darwin later reinterpreted this maxim through an evolutionary lens, arguing that it implies something more than resemblance—namely, "propinquity of descent" (Darwin 1859). In modern terms, characters help us recognize taxa, but they should not be conflated with the taxa themselves. While Linnaeus and Darwin had morphological characters in mind, the same caveat applies to molecular characters and the resulting topologies. A molecular tree may help identify a specific clade, but recognizing that clade and assigning it a Linnaean rank are different steps that require separate decisions. Topology can provide evidence that a candidate group is genealogically cohesive, but the further question of what Linnaean rank, if any, that group should receive requires additional judgment about diagnosability, stability, and utility. The central issue is therefore not whether phylogenetic evidence is useful in classification, nor whether monophyly is desirable when robustly supported. The issue is whether topology alone provides an adequate basis for ranked higher classification. We argue that it does not, and that the coronuloid case illustrates why: the nodes motivating rank changes prove sensitive to marker composition, taxon sampling, and analytical choices. As such, they cannot provide a foundation robust enough for redrawing a ranked classification.

Higher classification should therefore be treated as an interpretive step informed by phylogeny but evaluated against broader classificatory criteria —namely, diagnosability, stability, utility, and monophyly. Diagnosability requires that taxa convey biological information rather than absorb discordant forms under increasingly elastic definitions. Stability requires higher classification not to be repeatedly reshaped by unstable nodes, sparse taxon sampling, or the next incremental exercise in phylogeny. Utility requires that ranked taxa remain informative for the broader biological community, including paleontologists, ecologists, conservation biologists, educators, and policy makers. Monophyly remains fundamental because shared ancestry is a major source of systematic information, but should not function as an automatic command to redraw ranks whenever a preferred topology changes. These considerations support a more conservative working classification of Coronuloidea, which should continue to denote the epibiotic radiation of turtle and whale barnacles, whereas Tetraclitoidea should be retained for the tetraclitid–bathylasmatid–austrobalanid assemblage, comprising primarily rock-dwelling and deep-sea forms. Within the coronuloids, family-level changes should be approached cautiously unless they are supported not only by topology but also by explicit rediagnosis and a clear gain in information content. This is not an argument against phylogeny—rather it is an argument about how phylogenetic evidence should be translated into ranked taxonomy.

This matters because the same topology-driven changes that increase apparent conformity to monophyly can also reduce the diagnostic and comparative value of historically meaningful names such as Platylepadidae and Coronuloidea. Classificatory stability is therefore not merely a conservative preference but part of the infrastructure of biological communication. Similar concerns have recently been emphasized for biological nomenclature as a whole, with stability and universality being recognized as essential for unambiguous scientific communication (Jiménez- Mejías *et al*. 2024). Future total-evidence analyses integrating expanded molecular sampling, formal morphological matrices, multiple species of problematic genera, adequate representation of fossil taxa, and explicit comparisons with alternative classifications may well support a different result.

Until such evidence exists, the narrower, traditional concept of Coronuloidea and the retention of Platylepadidae provide the most informative classificatory scaffold.

### Beyond barnacles: what should higher classification accomplish?

The superfamily Coronuloidea provides an unusually strong case study because morphology, ecology, molecular phylogeny and the fossil record all bear directly on the same higher-classification problem. In this group, topology-driven classification has produced a broadened superfamily and revised family concepts that are more monophyly-oriented but less diagnosable, less biologically coherent, and less useful overall. That outcome supports a general conclusion: phylogeny is one relevant source of information for taxonomy, but it is not taxonomy itself; higher classification should take topology into account, but should not be determined by topology alone. Darwin was surely prescient in recognizing that classification must be grounded in descent (Darwin 1859), but the present case shows that drawing solely upon our own representations of descent cannot yield an unambiguous higher classification. The question for higher classification is therefore not simply whether a named group corresponds to a node, but whether that name continues to organize biological knowledge in a stable, diagnosable, and useful way.

### Taxonomic Account

In line with previous research by the first author, we retain *Tubicinella cheloniae* Monroe & Limpus, 1979 within *Tubicinella* rather than treating it as the type and sole species of *Chelolepas* Ross & Frick, 2007. Recognizing *Chelolepas* as a valid genus within Platylepadidae would necessitate assigning *Tubicinella* to Coronulidae. Definitive placements of these forms await future phylogenetic analyses and comprehensive morphological comparisons including *Tubicinella major* (the type species of *Tubicinella*), *T. cheloniae*, and the recently described fossil species †*Tubicinella nodai* Karasawa & Kobayashi, 2026.

Superfamily Coronuloidea Leach, 1817

Diagnosis (modified after Collareta *et al*. 2022): Balanomorphs with opercular plates occupying substantially less than the whole orifice area, ranging from weakly articulated to disarticulated, variously reduced or completely absent; wall either eight- or six-plated; parietes often with internal and/or external longitudinal parietal canals; external canals developed in-between external (secondary) T-shaped flanges, sometimes vestigial; basis membranous.

Family †Emersoniidae Ross in Ross & Newman, 1967

Diagnosis (modified after Harzhauser *et al*. 2011): Wall presumably six-plated (rostrum and rostrolatera fused to form a compound rostrum lacking any longitudinal sulcus, midrib and/or tooth), traversed by vertical and horizontal lamellae to form box-like cells.

genus †*Emersonius* Ross, 1967 (Eocene)

Family Chelonibiidae Pilsbry, 1916

Diagnosis (modified after Newman & Ross 1976): Wall eight- or six-plated, each plate lacking any longitudinal sulcus, midrib and/or tooth; opercular plates weakly articulated; terga well developed; one row of confluent internal longitudinal parietal canals formed between primary and secondary outer laminae.

genus *Chelonibia* Leach, 1817 (Miocene–Recent)

genus †*Protochelonibia* Harzhauser & Newman, 2011 (Oligocene–Pliocene)

Family Coronulidae Leach, 1817

Diagnosis (modified after Newman & Ross 1976): Wall six-plated, each plate lacking any longitudinal sulcus, midrib and/or tooth; two orders of longitudinal parietal canals well developed between primary and secondary T-shaped flanges; terga vestigial (opercular plates lacking in *Xenobalanus*).

genus †*Cetolepas* Zullo, 1969 (Pliocene)

genus *Cetopirus* Ranzani, 1817 (Pleistocene–Recent)

genus *Coronula* Lamarck, 1802 (?Miocene–Recent)

genus *Cryptolepas* Dall, 1872 (Pleistocene–Recent)

genus *Xenobalanus* Steenstrup, 1852

Family Platylepadidae Newman et Ross, 1976

Diagnosis (modified after Perreault *et al*. 2025): Wall six-plated (except for the obviously eight- plated juveniles of *Calyptolepas*); plates relatively thin; parietes each with one or more midribs and/or teeth (except in *Tubicinella*); two orders of longitudinal parietal canals often well developed between primary and secondary T-shaped flanges; terga moderately well developed.

genus †*Alabamalepas* Perreault et al., 2025 (Oligocene)

genus *Platylepas* Gray, 1825 (Pleistocene–Recent)

genus *Cylindrolepas* Pilsbry, 1916

genus *Calyptolepas* Frick et al., 2010

genus *Stomatolepas* Pilsbry, 1910

genus *Stephanolepas* Fischer, 1886

genus *Tubicinella* Lamarck, 1802 (Pleistocene–Recent)

## ACKNOWLEDGEMENTS

ChatGPT (OpenAI) and Gemini (Google LLC) were used to refine the English language and grammar of this manuscript.

## CONFLICT OF INTEREST

The authors declare no conflict of interest.

## DATA AVAILABILITY

The alignments, concatenated datasets, tree files, consensus tree files, and IQ-TREE output files supporting this study have been deposited in the Dryad Digital Repository: https://doi.org/10.5061/dryad.ngf1vhj96.

## AUTHOR CONTRIBUTION

RH conceived and designed the study; assembled and curated the molecular dataset; performed the phylogenetic analyses; examined the specimens; documented and interpreted the morphological characters; prepared the figures; and wrote the first draft of the manuscript. AC contributed additional expertise on morphology, fossil taxa, and the paleontological aspects of coronuloid classification. Both authors contributed to the taxonomic discussion, revised the manuscript critically for intellectual content, and approved the final version.

## FUNDING

This study was supported in part by the KAKENHI grant (JP19K04683 and JP26K15248 to RH) from the Japan Society for the Promotion of Science.

## SUPPORTING INFORMATION

Additional Supporting Information may be found in the online version of this article at the publisher’s web-site:

**Figure S1.**
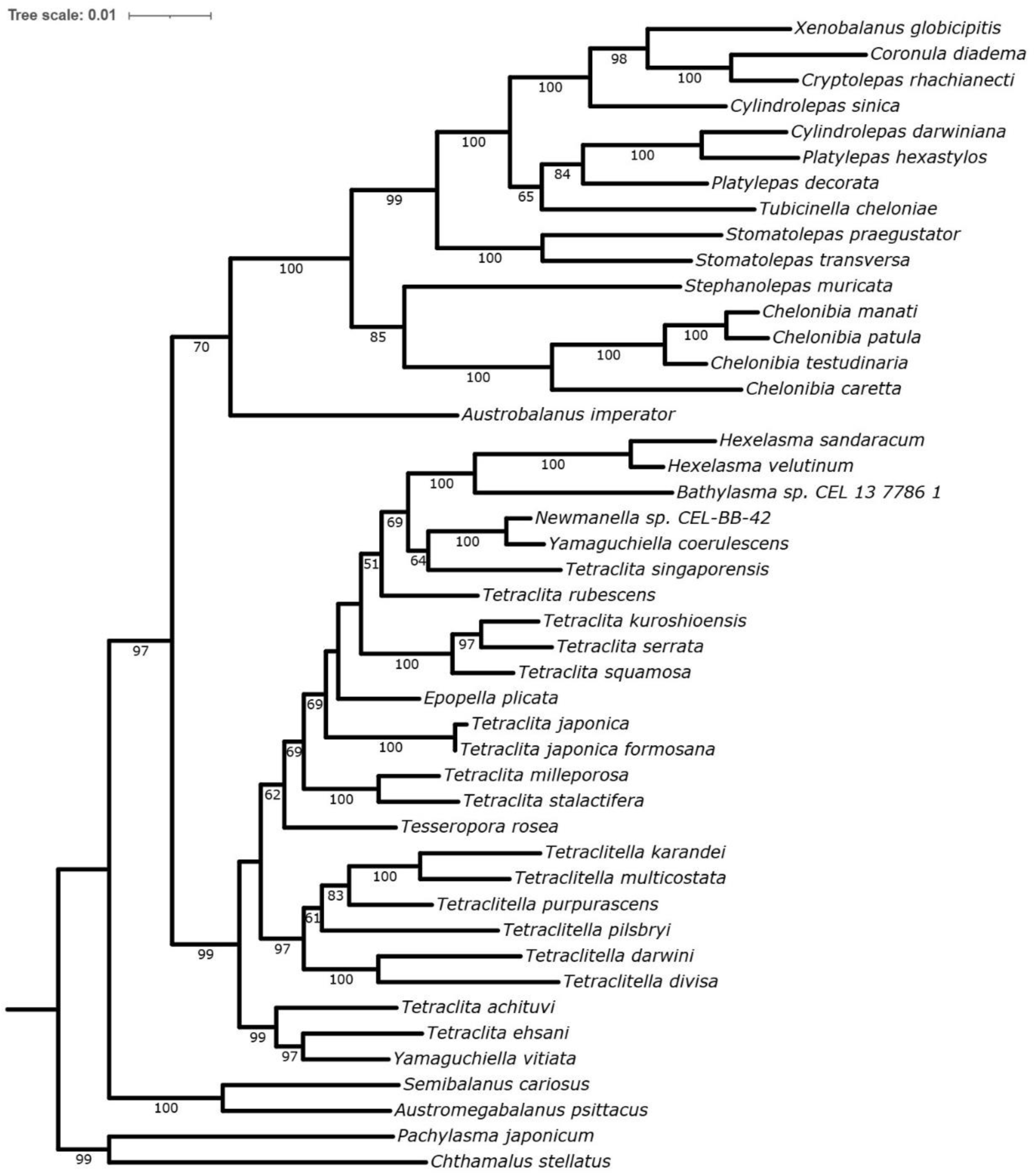
Maximum-likelihood phylogeny inferred from the concatenated four-marker dataset (12S, 16S, 18S, and H3) for 45 operational taxonomic units (OTUs). Numbers at nodes indicate bootstrap support values. Scale bar indicates substitutions per site.

**Table S1.**
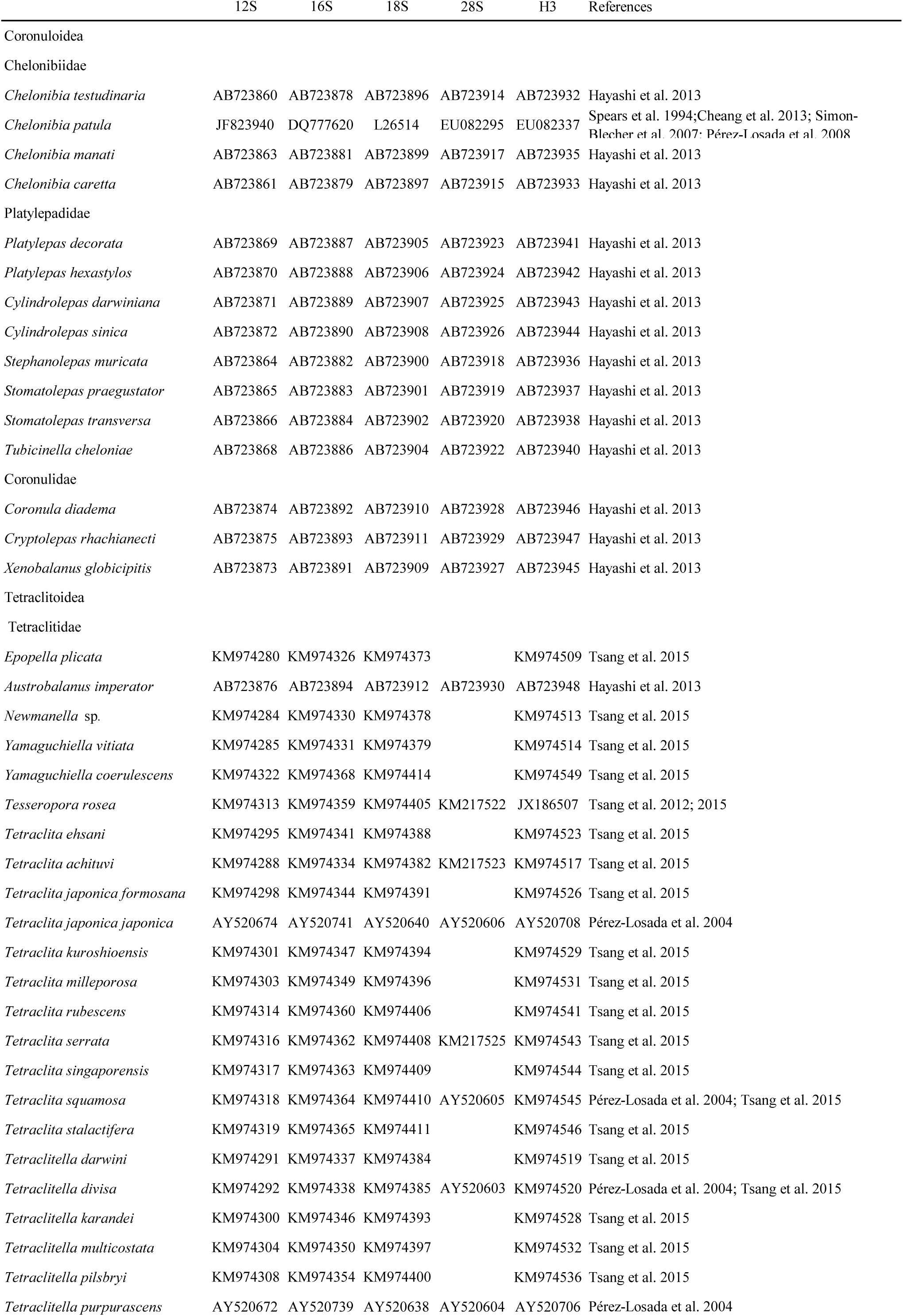
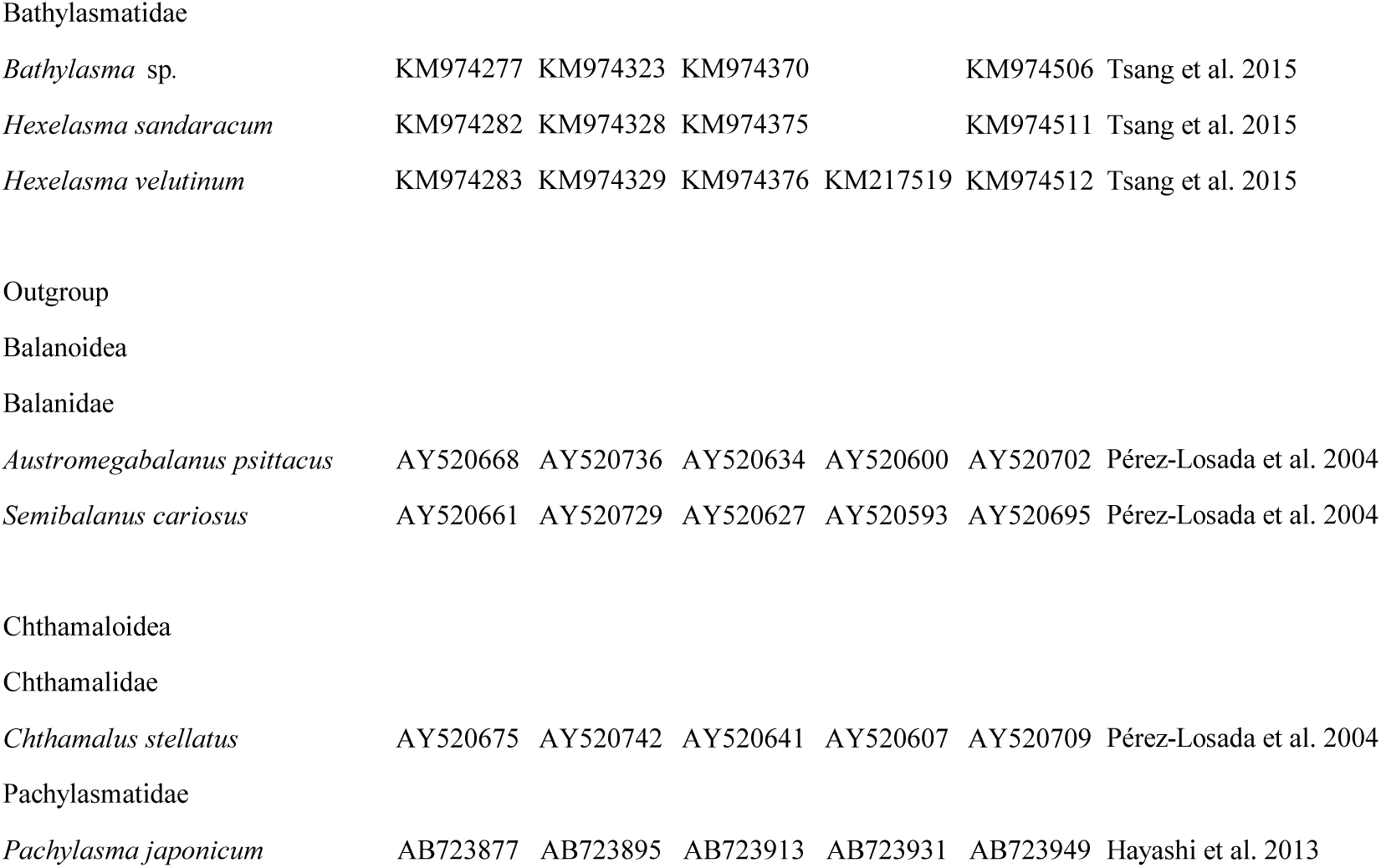
Information on the taxa included in the present study and GenBank accession numbers of the analyzed sequences. Note: Under the classification of Chan et al. (2021), Bathylasmatidae, Tetraclitidae, and Austrobalanidae are included within a broadened Coronuloidea. Here, these taxa are treated as part of the traditional tetraclitid–bathylasmatid–austrobalanid assemblage for comparative purposes.

## Notes

### Competing Interest Statement

The authors have declared no competing interest.

## REFERENCES

1. Ashlock PD. 1979. An evolutionary systematist’s view of classification. Systematic Zoology 28: 441–450.

2. Bigelow RS. 1958. Classification and phylogeny. Systematic Zoology 7: 49–59.

3. Bock WJ. 1973. Philosophical foundations of classical evolutionary classification. Systematic Zoology 22: 375– 392.

4. Brummitt RK. 1997. Taxonomy versus cladonomy, a fundamental controversy in biological systematics. Taxon 46: 723–734.

5. Chan BKK, Dreyer N, Gale AS, Glenner H, Ewers-Saucedo C, Pérez-Losada M, Kolbasov GA, Crandall KA, Høeg JT. 2021. The evolutionary diversity of barnacles, with an updated classification of fossil and living forms. Zoological Journal of the Linnean Society 193: 789–846.

6. Cheang CC, Tsang LM, Chu KH, Cheng IJ, Chan BKK. 2013. Host-specific phenotypic plasticity of the turtle barnacle *Chelonibia testudinaria*: a widespread generalist rather than a specialist. PLOS ONE 8: e57592.

7. Collareta A, Newman WA, Bosio G, Coletti G. 2022. A new chelonibiid from the Miocene of Zanzibar (Eastern Africa) sheds light on the evolution of shell architecture in turtle and whale barnacles (Cirripedia: Coronuloidea). Integrative Zoology 17: 24–43.

8. Dall WH. 1872. On the parasites of the cetaceans of the N.W. coast of America, with descriptions of new forms. Proceedings of the California Academy of Sciences 4: 299–301.

9. Darwin C. 1854. A monograph on the Subclass Cirripedia, with figures of all the species. The Balanidae, the Verrucidae, etc. London: The Ray Society.

10. Darwin C. 1859. On the origin of species by means of natural selection. London: John Murray.

11. Fischer P. 1886. Description d’un nouveau genre de Cirrhipèdes (Stephanolepas) parasite des tortues marines. Actes de la Société Linnéenne de Bordeaux 4: 193–196.

12. Franz NM. 2005. On the lack of good scientific reasons for the growing phylogeny/classification gap. Cladistics 21: 495–500.

13. Frick MG, Zardus JD, Lazo-Wasem EA. 2010. A new coronuloid barnacle subfamily, genus and species from cheloniid sea turtles. Bulletin of the Peabody Museum of Natural History 51: 169–177.

14. Gray JE. 1825. A synopsis of the genera of Cirripedes arranged in natural families, with a description of some new species. Annals of Philosophy, new series 10: 97–107.

15. Harzhauser M, Newman WA, Grunert P. 2011. A new Early Miocene barnacle lineage and the roots of sea-turtle fouling Chelonibiidae (Cirripedia, Balanomorpha). Journal of Systematic Palaeontology 9: 473–480.

16. Hayashi R. 2012. Atlas of the barnacles on marine vertebrates in Japanese waters including taxonomic review of superfamily Coronuloidea (Cirripedia: Thoracica). Journal of the Marine Biological Association of the United Kingdom 92: 107–127.

17. Hayashi R. 2013. A checklist of the turtle and whale barnacles (Cirripedia: Thoracica: Coronuloidea). Journal of the Marine Biological Association of the United Kingdom 93: 143–182.

18. Hayashi R, Chan BKK, Simon-Blecher N, Watanabe H, Guy-Haim T, Yonezawa T, Levy Y, Shuto T, Achituv Y. 2013. Phylogenetic position and evolutionary history of the turtle and whale barnacles (Cirripedia: Balanomorpha: Coronuloidea). Molecular Phylogenetics and Evolution 67: 9–14.

19. Jiménez-Mejías P, Manzano S, Gowda V. et al. 2024. Protecting stable biological nomenclatural systems enables universal communication: A collective international appeal. BioScience 74: 467–472.

20. Karasawa H, Kobayashi N. 2026. Some new records for cirripedes from the Pliocene–Pleistocene of Southwest Japan, with two new species of Balanomorpha. Bulletin of the Mizunami Fossil Museum 53: 9–21.

21. Katoh K, Rozewicki J, Yamada KD. 2019. MAFFT online service: Multiple sequence alignment, interactive sequence choice and visualization. Briefings in Bioinformatics 20: 1160–1166.

22. Kuntner M, Čandek K, Gregorič M, Turk E, Hamilton CA, Chamberland L, Starrett J, Cheng R.-C, Coddington JA, Agnarsson I, Bond JE. 2023. Increasing information content and diagnosability in family-level classifications. Systematic Biology 72: 964–971.

23. Lamarck JB. 1802. Mémoire sur la Tubicinelle. Annales du Muséum National d’Histoire Naturelle 1: 461–464.

24. Leach WE. 1817. Distribution systématique de la classe Cirripèdes. Journal de Physique, de Chimie, d’Histoire Naturelle et des Arts 85: 67–69.

25. Linnaeus C. 1751. Philosophia botanica. Stockholmiae: Apud Godofr. Kiesewetter.

26. Monroe R. 1981. Studies in the Coronulidae (Cirripedia). Memoirs of the Queensland Museum 20: 237–247.

27. Monroe R, Limpus CJ. 1979. Barnacles on turtles in Queensland waters with descriptions of three new species. Memoirs of the Queensland Museum 19: 197–223.

28. Newman WA. 1996. Sous-Classe des Cirripèdes (Cirripedia Burmeister, 1834) Super-Ordres des Thoraciques et des Acrothoraciques (Thoracica Darwin, 1854, Acrothoracica Gruvel, 1905). In Forest J. ed. Traité de zoologie. Anatomie, systématique, biologie. Paris: Masson, 453–540.

29. Newman WA, Ross A. 1976. Revision of the balanomorph barnacles: including a catalog of the species. Memoirs of the San Diego Society of Natural History 9: 1–108.

30. Pérez-Losada M, Høeg JT, Crandall KA. 2004. Unraveling the evolutionary radiation of the thoracican barnacles using molecular and morphological evidence: a comparison of several divergence time estimation approaches. Systematic Biology 53: 278–298.

31. Pérez-Losada M, Harp M, Høeg JT, Achituv Y, Jones D, Watanabe H, Crandall KA. 2008. The tempo and mode of barnacle evolution. Molecular Phylogenetics and Evolution 46: 328–346.

32. Perreault RT, Collareta A, Buckeridge JS. 2025. New fossils from the Oligocene of the southeastern U.S.A. evoke an ancient origin for the platylepadid turtle barnacles (Thoracica, Coronuloidea). Neues Jahrbuch für Geologie und Paläontologie - Abhandlungen 315: 57–66.

33. Pilsbry HA. 1910. *Stomatolepas*, a barnacle commensal in the throat of the loggerhead turtle. The American Naturalist 44: 304–306.

34. Pilsbry HA. 1916. The sessile barnacles (Cirripedia) contained in the collections of the U.S. National Museum; including a monograph of the American species. Bulletin of the United States National Museum 93: 1–366.

35. Ranzani C. 1818. Osservazioni su i Balanidi. Opuscoli Scientifici 2: 63–93.

36. Ross A, Frick MG. 2007. From Hendrickson (1958) to Monroe & Limpus (1979) and beyond: An evaluation of the turtle barnacle *Tubicinella cheloniae*. Marine Turtle Newsletter 118: 2–5.

37. Ross A, Newman WA. 1967. Eocene Balanidae of Florida, including a new genus and species with a unique plan of “Turtle-Barnacle” organization. American Museum Novitates 2288: 2–21.

38. Simon-Blecher N, Huchon D, Achituv Y. 2007. Phylogeny of coral-inhabiting barnacles (Cirripedia; Thoracica; Pyrgomatidae) based on 12S, 16S and 18S rDNA analysis. Molecular Phylogenetics and Evolution 44: 1333–1341.

39. Spears T, Abele LG, Applegate MA. 1994. Phylogenetic study of cirripedes and selected relatives (Thecostraca) based on 18S rDNA sequence analysis. Journal of Crustacean Biology 14: 641–656.

40. Steenstrup JJS. 1852. Om *Xenobalanus globicipitis*, en ny Cirriped-slaegt af Coronula familien. Videnskabelige Meddelelser fra den Naturhistoriske Forening i Kjøbenhavn 1852: 62–64.

41. Trifinopoulos J, Nguyen LT, von Haeseler A, Minh BQ. 2016. W-IQ-TREE: a fast online phylogenetic tool for maximum likelihood analysis. Nucleic Acids Research 44: W232–W235.

42. Tsang LM, Achituv Y, Chu KH, Chan BKK. 2012. Zoogeography of intertidal communities in the West Indian Ocean as determined by ocean circulation systems: patterns from the *Tetraclita* barnacles. PLOS ONE 7: e45120.

43. Tsang LM, Chu KH, Achituv Y, Chan BKK. 2015. Molecular phylogeny of the acorn barnacle family Tetraclitidae (Cirripedia: Balanomorpha: Tetraclitoidea): validity of shell morphology and arthropodal characteristics in the systematics of tetraclitid barnacles. Molecular Phylogenetics and Evolution 82: 324–329.

44. Vences M, Guayasamin JM, Miralles A, De La Riva I. 2013. To name or not to name: criteria to promote economy of change in supraspecific Linnean classification schemes. Zootaxa 3636: 201–244.

45. Wilkerson RC, Linton YM, Fonseca DM, Schultz TR, Price DC, Strickman DA. 2015. Making mosquito taxonomy useful: a stable classification of tribe Aedini that balances utility with current knowledge of evolutionary relationships. PLOS ONE 10: e0133602.

46. Zardus JD, Lake DT, Frick MG, Rawson PD. 2014. Deconstructing an assemblage of “turtle” barnacles: species assignments and fickle fidelity in *Chelonibia*. Marine Biology 161: 45–59.

47. Zullo VA. 1969. Thoracic Cirripedia of the San Diego Formation, San Diego County, California. Contributions in Science 159: 1–25.

